## Supplemental figures for "A task-general connectivity model reveals variation in convergence of cortical inputs to functional regions of the cerebellum"

### *Extended Data Figures*

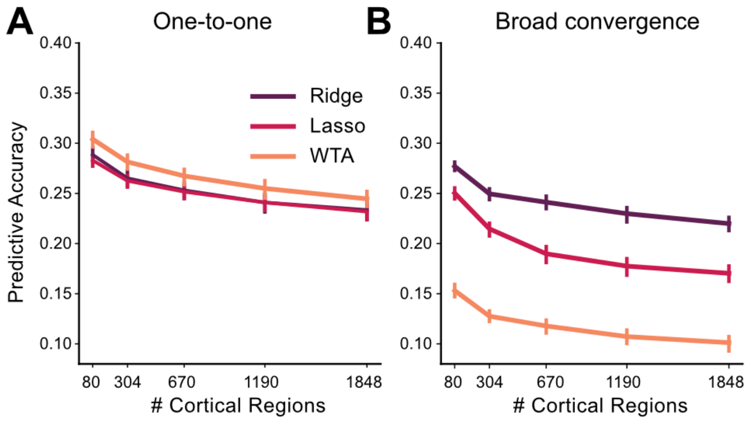


**Extended Data Fig. 1 | *Model recovery simulations demonstrate the ability to identify different forms of cortico-cerebellar connectivity.*** *Predictive accuracy for Ridge, Lasso, and WTA models trained and tested on simulated data generated using (**A**) one-to-one connectivity with each cerebellar voxel connected only to one randomly selected cortical region (**B**) broad convergence with connectivity weights being drawn from a Gaussian distribution. Data were simulated using the observed cortical activity for task set A and B for training and test data, respectively. The signal-to-noise level for the cerebellar data was matched to the empirical data. As expected, the WTA model provided the best prediction for the one-to-one architecture and the Ridge model provided the best prediction for the convergent architecture.*

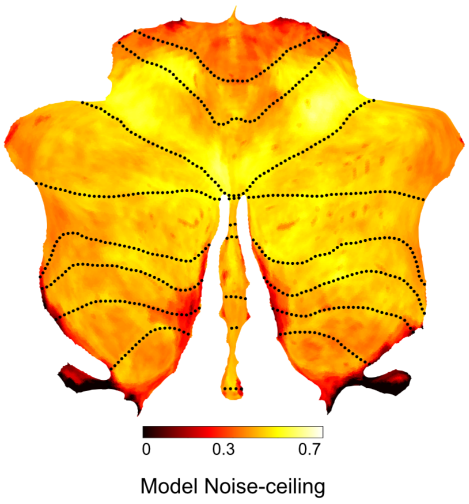


**Extended Data Fig. 2 | *Noise ceiling for Ridge model.*** *Expected predictive accuracy assuming that the fitted Ridge model (1848 parcels) reflects the true cortico-cerebellar connectivity. The noise ceiling takes into account the reliability of the cerebellar data, as well as the reliability of the prediction based on the measured cortical data.*

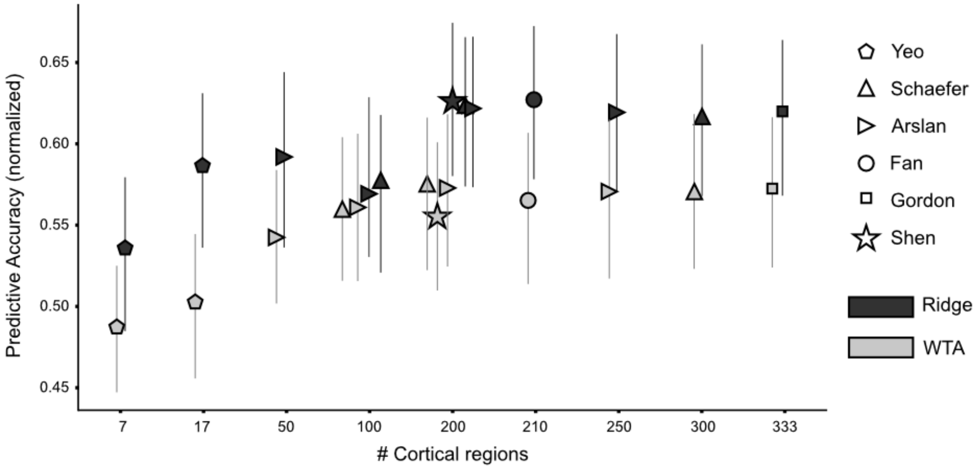


**Extended Data Fig. 3 | *Predictive accuracy for Ridge and WTA models using functional cortical parcellations*** *([25, 47, 48, 49, 50, 51], denoted by first author)***.** *Predictive performance is normalized to the noise ceiling. The number of cortical parcels varied from 7 to 330 regions. All evaluated parcellations are available at [github.com/DiedrichsenLab/fs_LR_32](https://github.com/DiedrichsenLab/fs_LR_32)[52].*

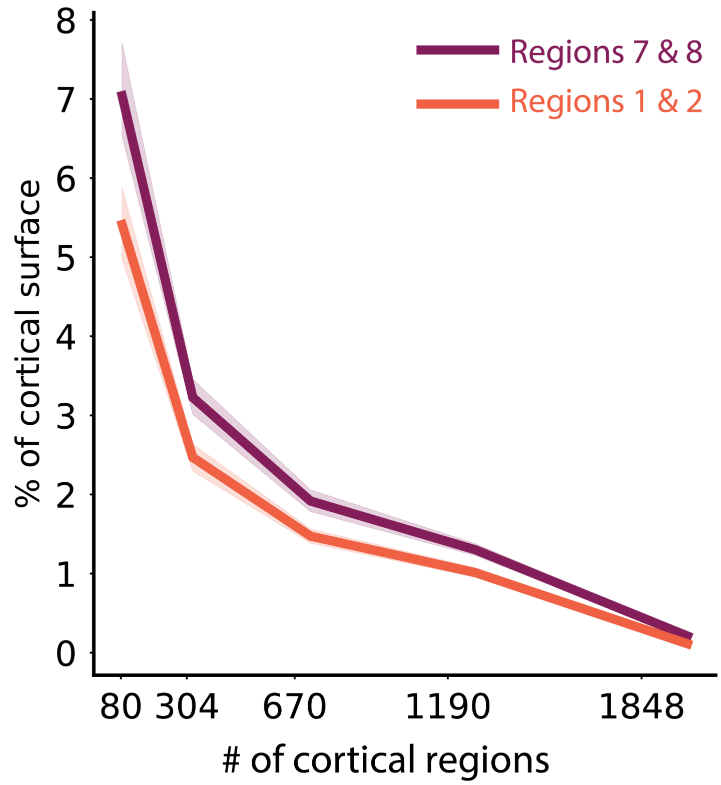


**Extended Data Fig. 4 | *Percentage of cortical surface across levels of granularity for Lasso regression model.*** *Calculation is performed as in Figure 3b. The results are averaged across the two hand regions of the cerebellum (MDTB functional parcellation, regions 1 and 2), and regions related to narrative and word comprehension (regions 7 and 8).*

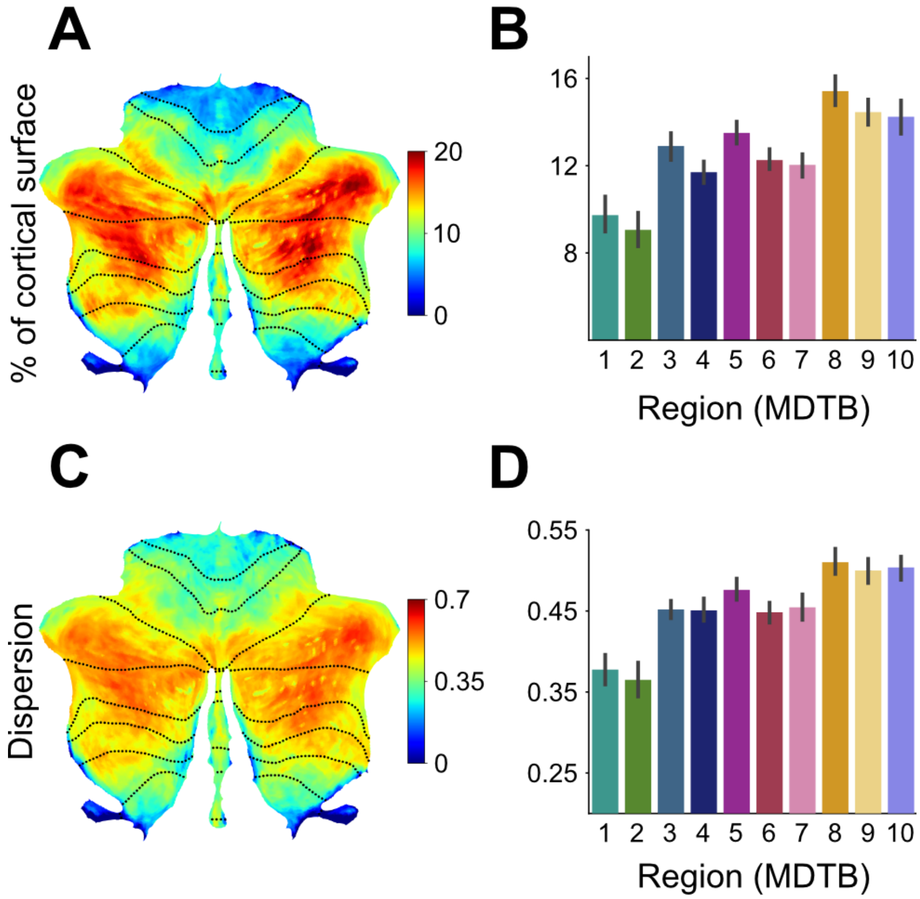


**Extended Data Fig. 5 | *Cortico-cerebellar convergence measures using the Ridge model.*** *A) Map of the cerebellum showing percentage of cortical parcels with coefficients for the Ridge model (n=80 parcels) above threshold (see methods). B) Percentage of parcels with weights above threshold for functional subregions of the cerebellum. C) Spherical dispersion of the connectivity weights on the cortical surface for each cerebellar voxel. D) Average cortical dispersion for each functional subregion of the cerebellum. Error bars indicate standard error of the mean across participants.*

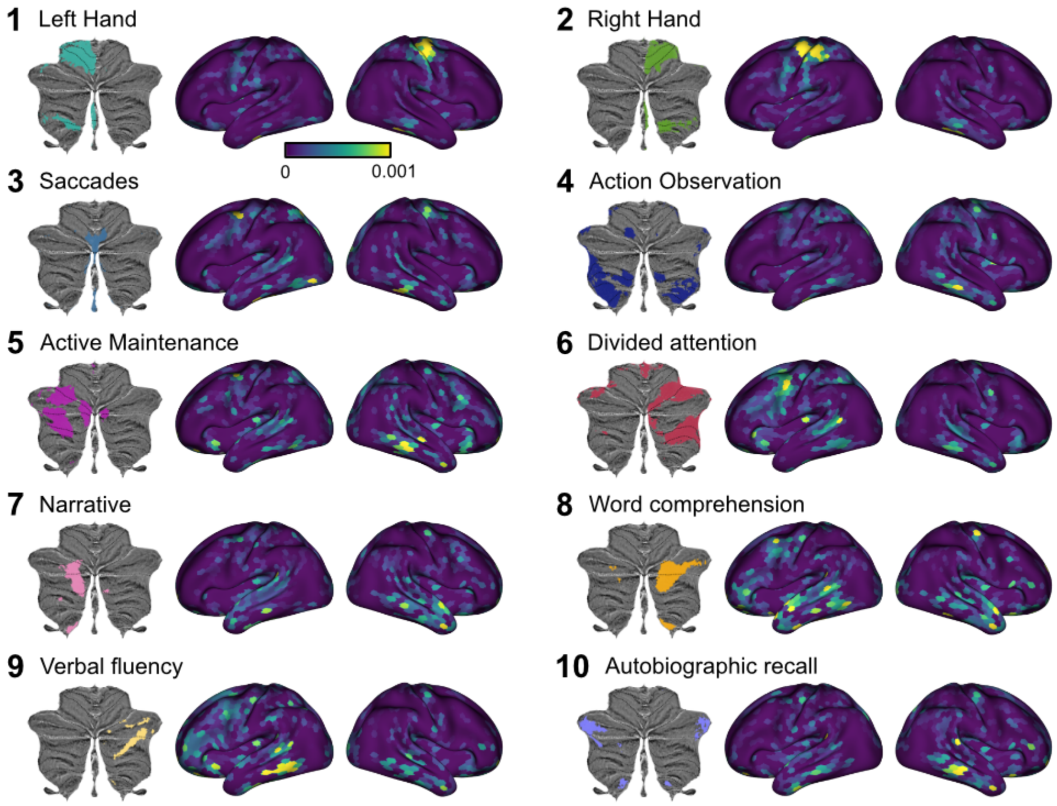


**Extended Data Fig. 6 | *Cortical connectivity weight maps for the Lasso model with 1848 cortical regions.*** *As in Figure 4, the results are averaged across individuals for each of the 10 functional regions defined on the MDTB data set, with each region denoted by the most important functional term. Results are averaged across participants and voxels within each cerebellar region. Regression weights are in arbitrary units.*

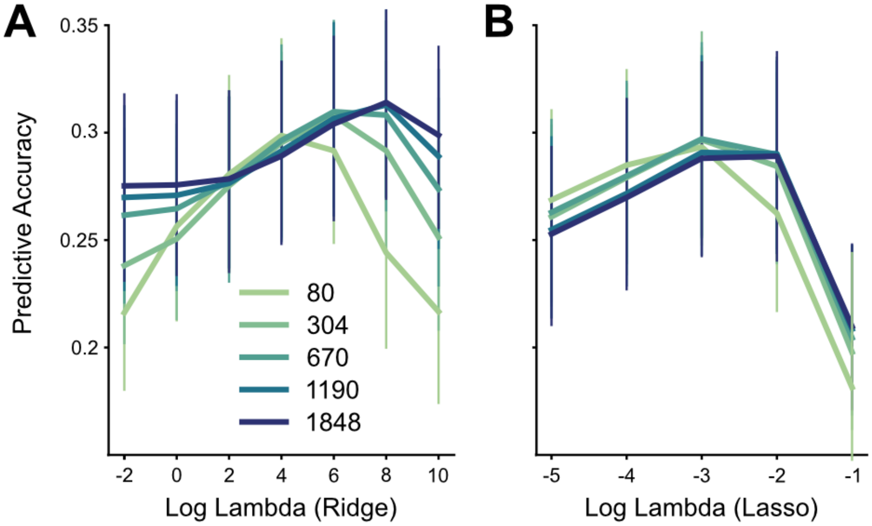


**Extended Data Fig. 7 | *Hyper-parameter tuning for connectivity models.*** *Predictive accuracy for Ridge (A) and Lasso (B) models using different regularization coefficients (log-lambda values) across five levels of granularity, and cross-validated over 4 folds of the training data. The WTA model did not use a hyper-parameter.*
